# Environmental circadian disruption re-programs liver circadian gene expression

**DOI:** 10.1101/2023.08.28.555175

**Authors:** Hao A. Duong, Kenkichi Baba, Jason P. DeBruyne, Alec J. Davidson, Christopher Ehlen, Michael Powell, Gianluca Tosini

**Affiliations:** Department of Pharmacology and Toxicology, Morehouse School of Medicine, Atlanta GA 30310; Neuroscience Institute, Morehouse School of Medicine, Atlanta GA 30310; Department of Microbiology, Biochemistry and Immunology, Morehouse School of Medicine, Atlanta GA 30310

## Abstract

Circadian gene expression is fundamental to the establishment and functions of the circadian clock, a cell-autonomous and evolutionary-conserved timing system. Yet, how it is affected by environmental-circadian disruption (ECD) such as shiftwork and jetlag, which impact millions of people worldwide, are ill-defined. Here, we provided the first comprehensive description of liver circadian gene expression under normal and after ECD conditions. We found that post-transcription and post-translation processes are dominant contributors to whole-cell or nuclear circadian proteome, respectively. Furthermore, rhythmicity of 64% transcriptome, 98% whole-cell proteome and 95% nuclear proteome is re-written by ECD. The re-writing, which is associated with changes of circadian cis-regulatory elements, RNA-processing and protein trafficking, diminishes circadian regulation of fat and carbohydrate metabolism and persists after one week of ECD-recovery.

**One-Sentence Summary:** Environmental-circadian disruption re-writes the circadian gene expression process which persists after one week of recovery.

## MAIN TEXT

The circadian clock, a cell-autonomous timing system, is driven by an evolutionary conserved transcription-translation feedback loop. It has evolved as a mechanism for organisms to synchronize daily cycles of internal biological rhythms with external environmental conditions by modulating the expression of thousands of clock-controlled genes(*1–4*). Under <u>e</u>nvironmental <u>c</u>ircadian <u>d</u>isruption (ECD) conditions such as jetlag, social jetlag and shiftwork, the natural harmonic alignment is disrupted, resulting in acceleration of neurological, cardiometabolic and immune disorders and cancers(*5–12*). This is a growing global health challenge, affecting hundreds of millions of people worldwide. In the United States and European countries, about 16% the workforce is subjected to a such condition via shiftwork, and the percentage is expectedly to be higher in developing countries. Yet how the circadian gene expression system is affected by ECD remains unclear. To address this question as well as illuminate its molecular underpinnings and functional implications, we systematically interrogated changes in the circadian gene expression process – encompassing transcription, translation and post-translation – at the - omics level in mouse livers after one-week ECD recovery(*13*) in comparison to standard circadian (STD) conditions (fig. S1).

### Integrated circadian gene expression

The circadian gene expression process had been intensively investigated, resulting in a thorough understanding of circadian transcription regulation from chromatin opening, cis-trans regulator interaction to transcript abundance(*14–16*). However, at the post-transcriptional and post-translational levels our understanding of the process is sparse(*17–20*). To fill this gap as well as inquire the state of gene expression under the STD control condition, we performed RNA-seq of whole-cell mRNAs and mass spectrometry analysis of whole-cell and nuclear proteins in the same set of mouse livers, which were entrained under STD and collected under circadian conditions (fig. S1).

#### Topography of circadian gene expression

Among 21061 quantified transcript time series, we found 5502 (26%) exhibit a rhythmic pattern of abundance with the peaks spreading throughout the cycle. Transcripts of all core clock genes including *Clock, Arntl, Per1/2/3, Cry1/2, Csnk1d/e* and *Nr1d1/2* were quantified in the RNA-seq. Their patterns of abundance were validated by RT-qPCR (Figs 1A-B, S2A) and are consistent with previous studies(*15, 21*). At the protein level, we quantified 7,314 circadian time series across whole-cell and nuclear compartments. Processing and results of mass-spec were validated in terms of enrichment of known mouse liver nuclear proteins, Pearson correlation between compartments, consistence of patterns of circadian protein abundance between mass-spec and published studies(*22, 23*) and enrichment of known circadian associated functions (figs. S2B-E). In the whole-cell circadian proteome, 285 (3.9%) were rhythmic. These proteins exhibit a bimodal distribution of peak abundance concentrating around CT7 and CT17. In the nuclear compartment, 678 (9.3%) were rhythmic with a distinct phase distribution from that of the whole-cell proteome, assuming a unimodal pattern centering around CT20 (Fig. 1B). These proteomes are enriched with cellular and molecular processes such as endosome transport, autophagy and response to hypoxia (fig. S2E). This is the first comprehensive and coherent description of daily abundance of whole-cell transcripts, whole-cell proteins and nuclear proteins in the same set of tissues under circadian condition.

**Fig. 1:**
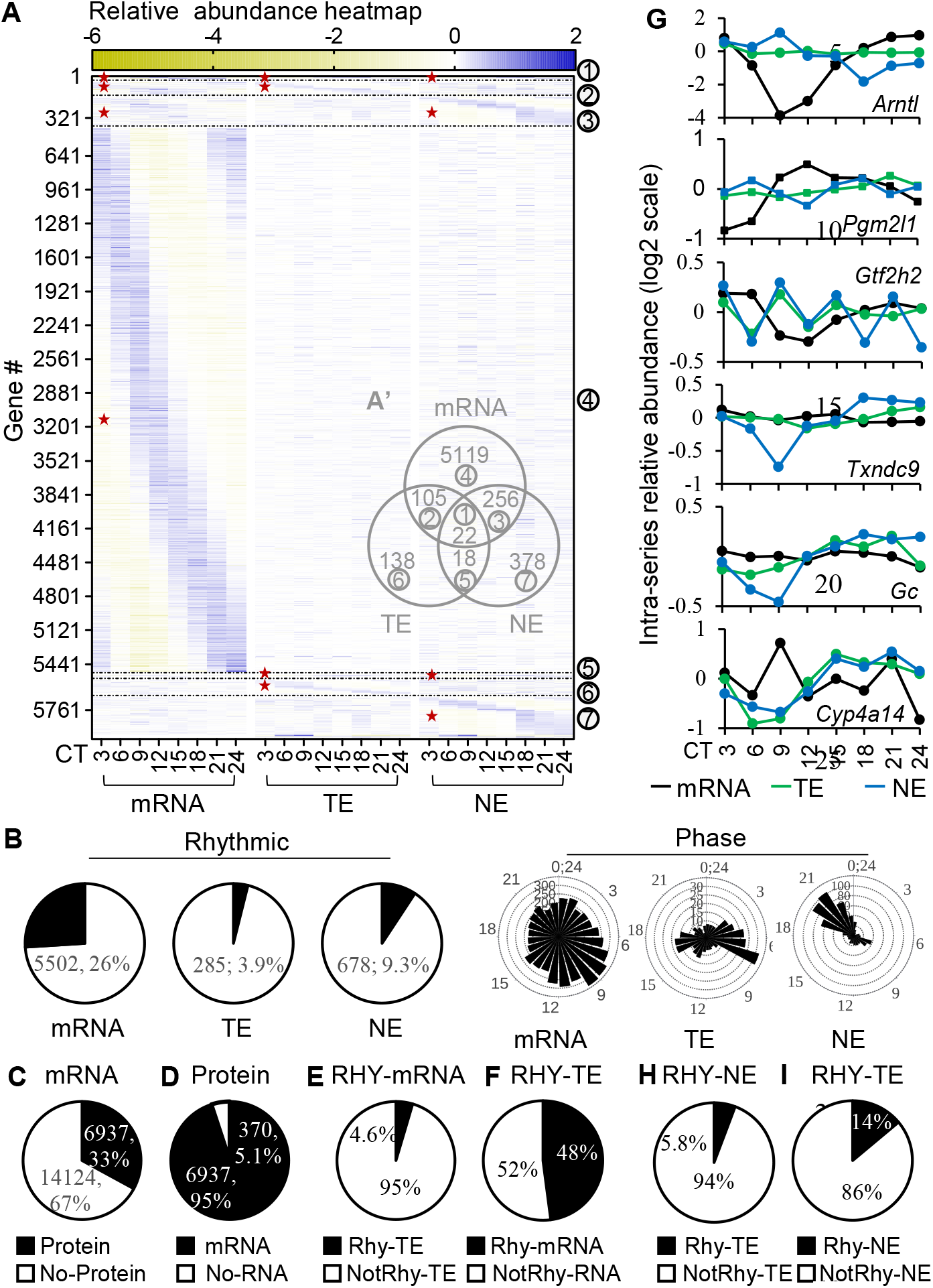
A topography of circadian gene expression in mouse liver. (**A, A’**) Relative-abundance heatmap (normalized to series average; log2 scale) of rhythmic transcripts (mRNA), rhythmic whole-cell proteins (TE) and rhythmic nuclear proteins (NE) (A), and their Venn diagram of rhythmic overlaps (A’). (**B**) Percentages of rhythmic time series and their phase distribution of transcripts, whole cell proteins and nuclear proteins. (**C-F**) Percentage of transcripts with or without proteins (C), proteins with or without transcripts (D), rhythmic transcripts with or without rhythmic whole-cell proteins, (E) rhythmic whole-cell proteins with or without rhythmic transcripts (F) in the 6937 sub-population. (**G**) Examples of differential transcript (transcription), whole-cell protein (translation) and nuclear (post-translation) circadian patterns. (**H-I**) Percentages of rhythmic whole-cell proteins with or without rhythmic nuclear proteins (H), or rhythmic nuclear proteins with or without rhythmic whole-cell proteins (I) in the 6937 sub-population. TE – total extract; NE – nuclear extract; RHY – rhythmic; “*”– rhythmic portion; 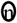 – corresponding groups between (A) and (A’).

#### Differential rhythmicity of transcripts and proteins

The rhythmicity of a transcript abundance has routinely been used to infer circadian biological function, but it is protein that mostly performs the gene function. We thus asked how faithfully transcript rhythmicity is recapitulated in protein rhythmicity. To minimize the effect of difference in detection levels between RNA-seq and Mass Spectrometry on the analysis, we firstly selected for genes of which transcript(s), whole-cell protein(s) and nuclear protein(s) were all quantified in the circadian transcriptome and circadian proteomes, and then examined their relationship in time and space. We found 6937 of such genes, constituting 33% all quantified transcripts and 95% all quantified proteins (Figs. 1C-D). Within this sub-population, less than 5% rhythmic transcripts associate with rhythmic whole-cell proteins and more importantly 52% rhythmic whole-cell proteins don’t associate with rhythmic transcripts (Figs. 1E-F). The percentages showed that most rhythmic transcripts don’t become rhythmic proteins and the majority of rhythmic proteins don’t directly originate from rhythmic transcripts. They also suggested that post-transcription contributes as much as transcription to circadian whole-cell protein rhythmicity. This is a higher percentage than previously reported of ∼20% in the same tissue under the same condition(*17*) and is close to ∼40% reported for the same tissue under diurnal condition(*24, 25*). Moreover, numerous genes play important roles in key biological processes such as transcription (*Arntl, Pgm2l1, Gtf2h2, Txndc9, Polr1d, Polr2c*) or metabolism (*Gc, Cyp4a14, Gpat4, Acot12)* exhibit differential circadian rhythmicity at transcript and protein levels (Fig 1G). These results indicate a profound dissociation in circadian rhythmicity of transcripts and proteins and thus warrant extra caution in inferring a circadian function from rhythmic transcripts at individual or -omics levels.

#### Post-translation underlies circadian nuclear proteom

Whole-cell protein abundance measures the collective amount of a protein in a cell but doesn’t carry any information on its distribution in sub-cellular compartments where cell functions are largely performed. To see how sub-cellular localizations, which are active and tightly regulated processes(*26*), affect circadian protein rhythmicity, thereby circadian outputs, we examined the overlap between whole-cell and nuclear circadian proteomes. 94% nuclear rhythmic proteins are not rhythmic at the whole cell level (Fig. 1H). The rhythmicity of these proteins must be acquired post-translationally as they are not rhythmic at the whole-cell level, making contribution from transcriptional, post-transcriptional and translational controls unlikely. Additionally, only 14% of whole-cell rhythmic proteins might be allocated to the rhythmicity in nuclear compartment (Fig. 1I). The remainders, 86%, have to be attributed to rhythmicity in other compartments, re-constituting their rhythmicity at the whole-cell level. These results indicate the circadian nuclear proteome mostly arises at the post-translational level and suggest the existence of compartmental specific circadian proteomes, a less studied area in the field(*27*), not only in the nucleus but also in other organelles.

### ECD re-writes the circadian transcriptome

After ECD we found 3445 (16%) transcripts exhibiting a circadian rhythmic pattern of abundance among 21061 quantified circadian time series. In response to ECD, 3118 STD rhythmic transcripts lose their rhythmicity (LOR_R_) while 1061 non-rhythmic transcripts acquired rhythmicity (GOR_R_), constituting 56.7% and 30.8% of all rhythmic transcripts under the same condition, respectively. There are 2384 STD rhythmic transcripts that retain rhythmicity (ROR_R_), regardless of whether there is a change in phase or amplitude of the oscillation. Each of these rhythmic populations displays a distributed pattern of peak abundance throughout the cycle (Fig 2A-C). Collectively, there are 6,563 rhythmic transcripts, of which 64% switched rhythmicity in response to ECD (Fig. 2D). The circadian transcriptome is thus re-written by ECD.

**Fig. 2:**
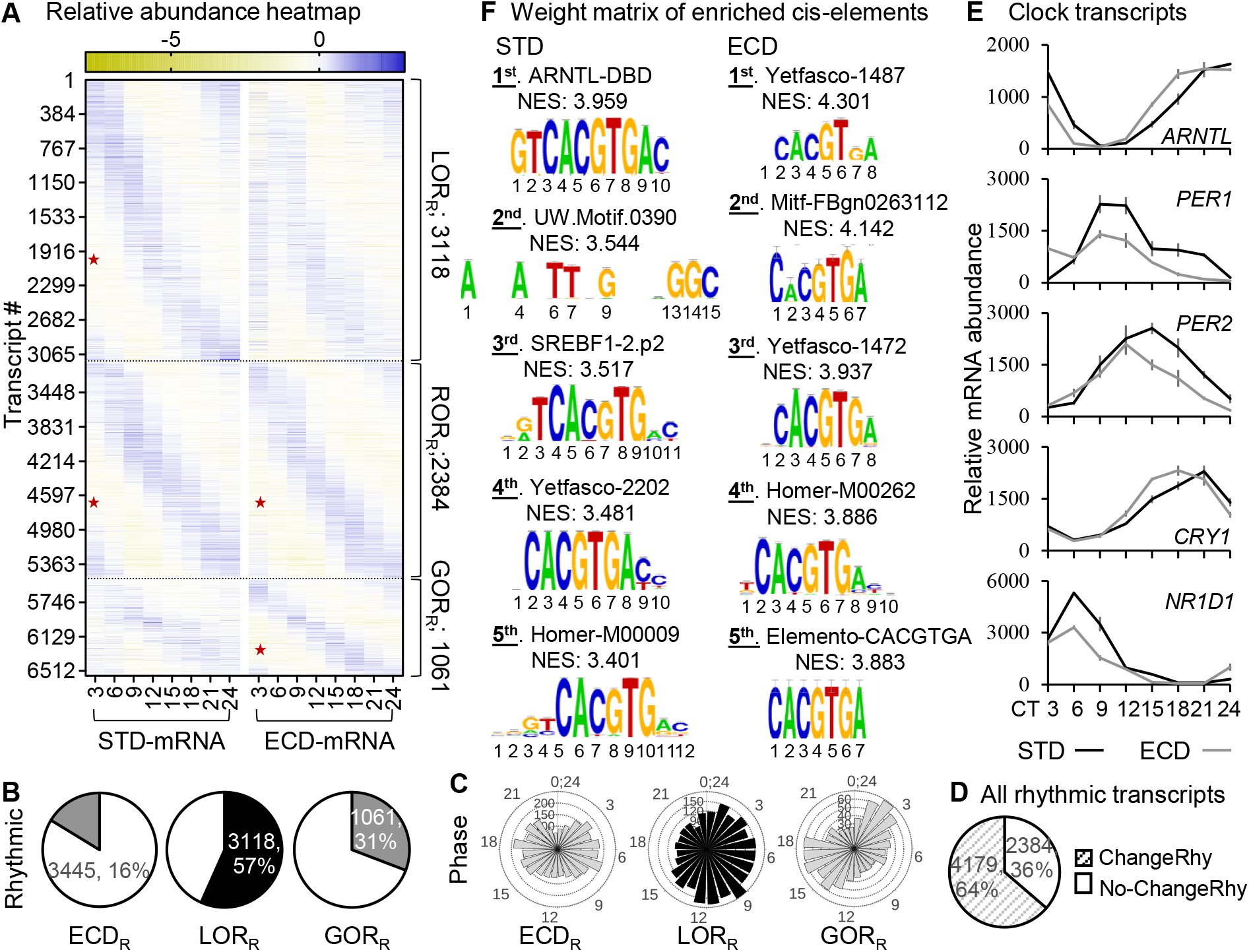
ECD re-writes rhythmicity of 4179 transcripts. (**A**) Relative-abundance heatmap (normalized to series average; log2 scale) of rhythmic transcripts in 21061 transcripts that were quantified under both standard (STD) and ECD conditions. (**B**) Percent rhythmic transcript time series and the corresponding phase distribution of transcripts that are either rhythmic under ECD, rhythmic under STD but not under ECD (LOR_R_), not rhythmic under STD but rhythmic under ECD (GOR_R_) or rhythmic under both STD and ECD (ROR_R_). (**C**) The total amount of transcripts that switch their rhythmicity in response to ECD. (**D**) Core clock transcripts that showed a change in their circadian pattern in response to ECD as quantified by RNA-seqs. (**E**) Weight matrix of enriched cis-elements within 20kb upstream of the either STD or ECD rhythmic transcripts ranked by Normalized Enrichment Score (NES); “*”– rhythmic portion.

Among core clock transcripts, the rhythmicity of *Arntl, Per1/2, Cry1* and *Nr1d1* showed a perturbation in phase or amplitude, but none exhibits a marked loss or gain of rhythmicity (Fig. 2E, fig. S3), suggesting potential contribution of factor(s) other than the core clock machinery to the re-writing. To seek clues for such regulator(s), we compared enrichment of cis regulatory elements within 20Kb upstream of transcription start sites in STD-rhythmic and ECD-rhythmic populations using iRegulon(*28*). At the top of the list for the STD-rhythmic population (5502 transcripts) is “ARNTL-DBD”, a motif for DNA binding domain of ARNTL(or BMAL1) which is a key transcription activator of the circadian feedback loop(*29*). Such enrichment is therefore expected and a validation of the analysis. Among the top 5 STD enriched elements, 4 harbor the E-box (CACGTG), a known dominant circadian cis-element(*30*). In ECD-rhythmic population, the highest enriched element is not a known circadian cis-element. E-box comes at 3^rd^ to 5^th^ positions. Thus, re-writing of circadian rhythmic transcripts associates with a reduction of BMAL1 binding concurrence with an increase in binding of a to-be-identified transcription factor(s) (Fig. 2F).

### Circadian whole-cell and nuclear proteomes are re-written by ECD

After ECD, we quantified 5317 protein time series across both whole-cell and nuclear compartment. At the whole-cell level, 909 (17%) series exhibit a circadian rhythmic pattern of abundance. This population displays a sharp bimodal distribution of peak abundance around 3 hours before anticipated dark (CT9) or dawn (CT21). Among 285 whole-cell STD rhythmic proteins, only 24 are also rhythmic after ECD (ROR_TE_). 261 proteins, or 92% of STD rhythmic proteins, lost their rhythmicity (LOR_TE_) while 885 proteins, which is about 3 times the number of STD rhythmic proteins, gained rhythmicity (GOR_TE_) in response to ECD. Collectively, ECD switched the rhythmicity of 1146 proteins, constituting 98% of all rhythmic proteins under both conditions at the whole-cell protein level. (Figs. 3A, C, E).

**Fig. 3:**
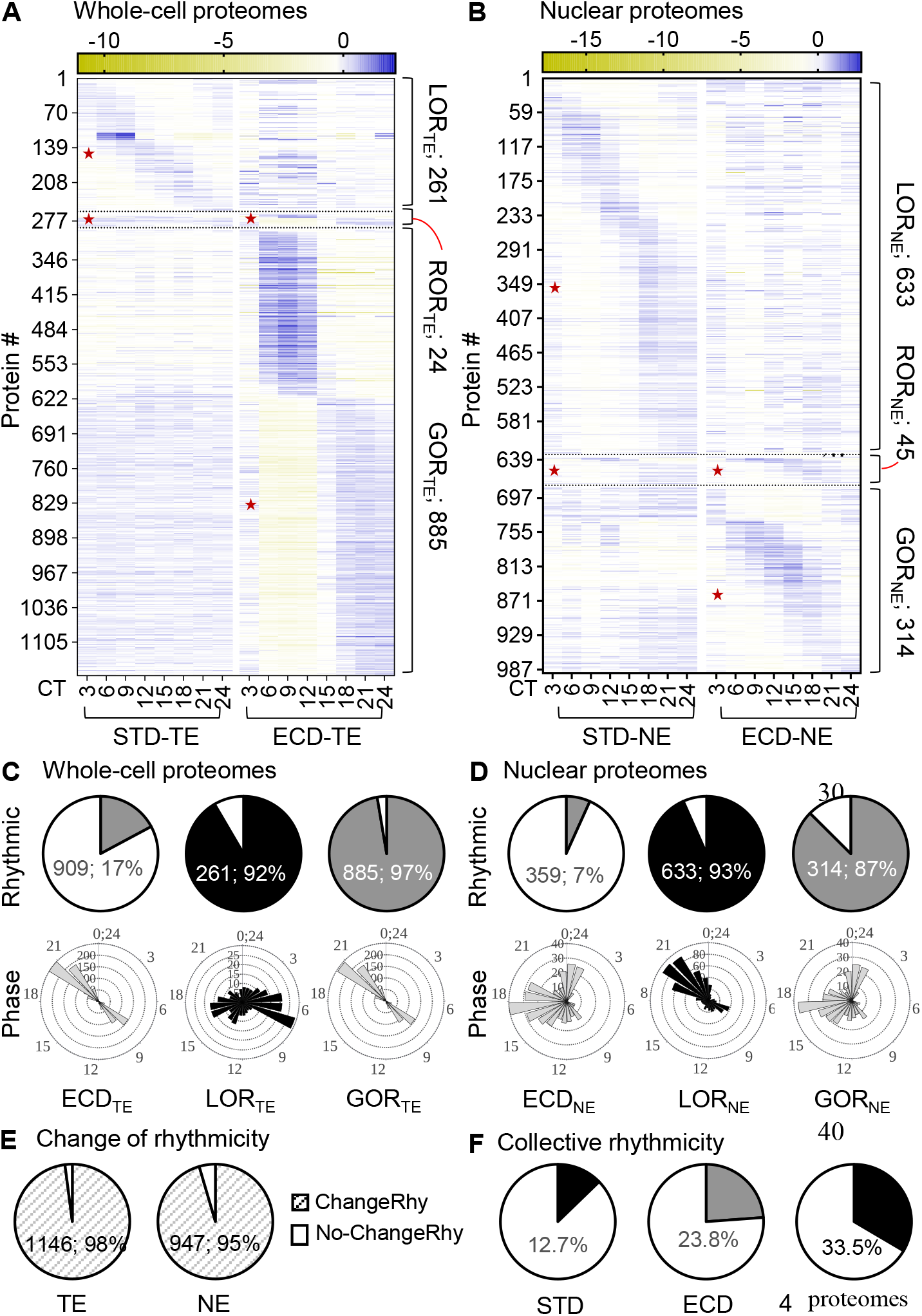
ECD re-writes rhythmicity of circadian whole-cell (TE) and nuclear (NE) proteomes. (**A-B**) Relative-abundance heatmap (normalized to series average; log2 scale) of rhythmic proteins in whole-cell (A) or nuclear (B) circadian proteomes under ECD in comparison with STD. (**C-D**) Percent rhythmic protein time series and the corresponding phase distribution of proteins that either loss rhythmicity (LOR), retain rhythmicity (ROR) or gain rhythmicity (GOR) in the whole-cell (C) or nuclear (D) circadian proteomes. (**E**) Percent of all proteins that change their rhythmicity at the whole-cell or nuclear compartment levels. (**F**) Estimate percentages of rhythmic proteins, collectively; “*”– rhythmic portion.

In the nuclear compartment, 359 (7%) protein time series are rhythmic with their phases distributed throughout a large part of the cycle. 633 of 678 STD rhythmic proteins (93.4%) lost their rhythmicity (LOR_NE_) while 314 proteins acquired rhythmicity (GOR_NE_) in response to ECD. There are 992 proteins, in total, that exhibit a rhythmic pattern of abundance in the nuclear compartment, of which 95% switched their rhythmicity after ECD. (Figs. 3B, D, E). Collectively, 12.7%, 23.8% and 33.5% proteins are rhythmic under STD, ECD or both conditions, respectively (Fig. 3F).

Such high magnitudes of change in rhythmicity at the transcript and protein levels were unexpected. To see if loosening the stringency for rhythmic calling would have a significant impact on the magnitudes, we re-performed the analysis using one of the highest recall (RAIN)(*31*) or most popular (JTK-Cycle)(*32*) algorithm. Both showed insignificant and mild changes in the magnitude of re-writing of the proteomes or transcriptome, respectively (fig. S4). These observations suggested that ECD re-writes the entire circadian gene expression system from transcription to post-translation.

The increase in protein rhythmicity at the whole-cell level, both quantity (from 3.9% to 17%) and synchrony (dispersed to discrete bimodal phase distribution), after ECD compared to STD is intriguing (Fig. 1B-TE vs Fig. 3C-ECD_TE_). The circadian clock has evolved as a mechanism to help organisms synchronize their internal biological processes with anticipated changes of daily environmental cycles. One would expect a lessening, rather than increasing, of protein rhythmicity under environmental circadian disruption. However, biological systems are composed of interactive networks, which help their resilience to external perturbations by redistributing, in part, the impact. Under a circadian disrupted condition, these networks are likely compromised, making their nodes more vulnerable and thereby weakening the resilience. From this perspective, ECD could lead to more protein rhythmicity in a synchronous manner, which was what we observed. In agreement with this interpretation, resilience of processes such as inflammation, glomerular and tubular injuries is compromised after ECD(*6, 33, 34*). We thus propose that the gain of rhythmicity reflects not a strengthening of the system but a losing of resilience to perturbations.

### Post-transcription and post-translation are dominant contributors to circadian proteomes under ECD

Given that post-transcription and post-translation are the major contributors to protein rhythmicity under STD, we asked how ECD would affect those contributions. We examined overlaps between the circadian transcriptome, circadian whole-cell proteome and circadian nuclear proteome after ECD as performed for STD. After ECD, we found 5028 transcripts of which whole-cell protein or nuclear protein were also quantified. Within the 5028 sub-population, 73% rhythmic whole-cell proteins don’t have rhythmic transcripts (Figs. 4 A-E). This percentage is substantially higher than the 52% under STD. In the nuclear protein rhythmic sub-population, 86% are not rhythmic at the whole-cell protein level (Figs. 4F-G). Thus, under ECD post-transcriptional and post-translational processes are dominant contributors to protein rhythmicity at both whole-cell and nuclear compartment levels.

**Fig. 4:**
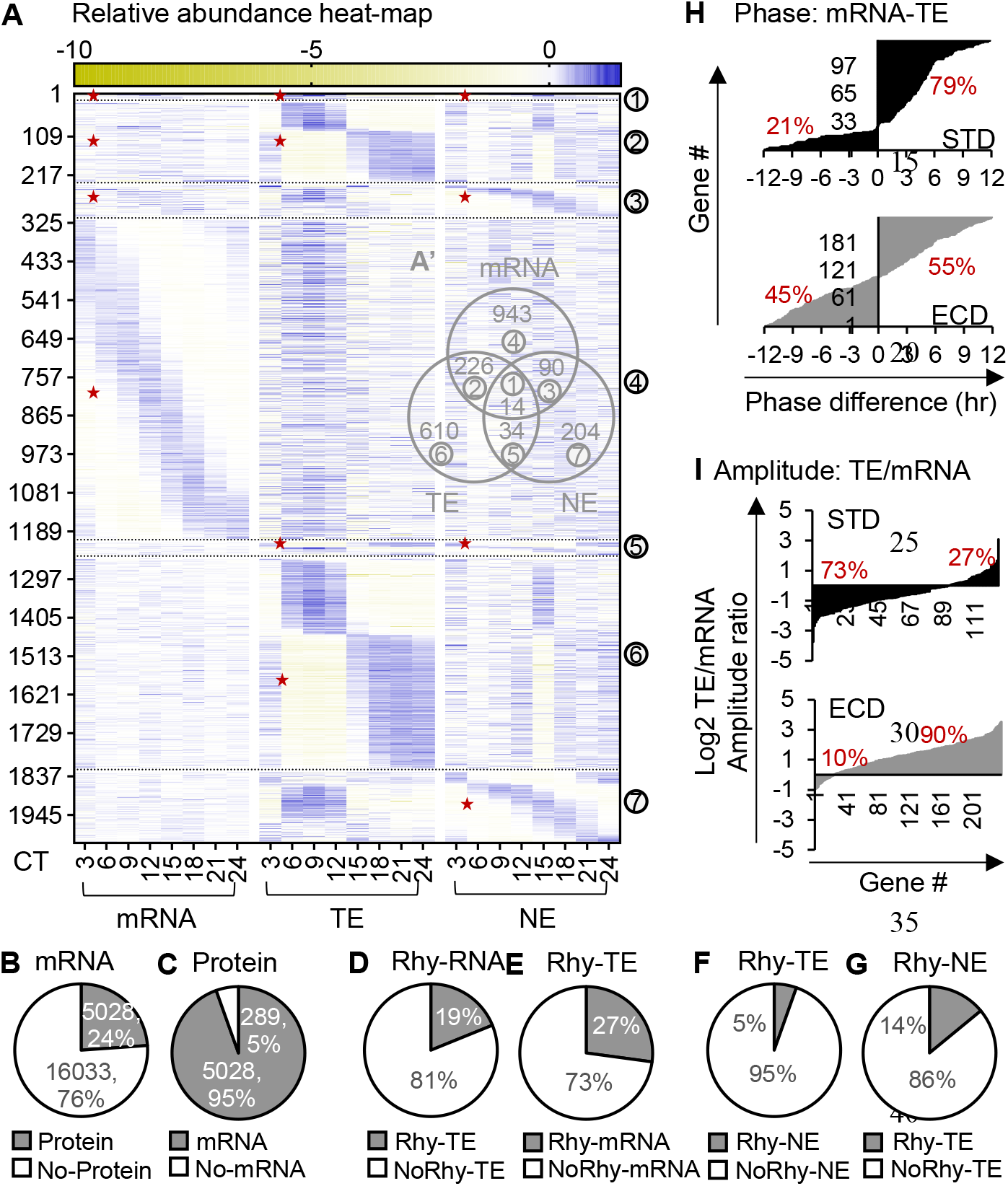
Contributions of post-transcription and post-translation in circadian gene expression under ECD. (**A**) Relative abundance heatmap (log2) of rhythmic transcripts, rhythmic whole-cell protein and rhythmic nuclear proteins in 5028 genes of which both transcript and protein were quantified under ECD. (**A’**) Venn diagram shows overlaps between each population shown in A. (**B-C**) Percent of transcripts with or without proteins and vice versa. (**D-E**) Percent of rhythmic transcripts with or without rhythmic whole-cell proteins and vice versa. (**F-G**) Percent of rhythmic whole-cell proteins with or without rhythmic nuclear proteins and vice versa. (**H**) Percent of genes with the phase of transcript leads or lags the phase of whole-cell protein under STD or ECD. (**I**) Percent of genes with the amplitude of whole-cell protein rhythms is lower or higher than that of the transcript rhythms under STD or ECD; “*”– rhythmic portion; 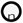 – corresponding groups between (A) and (A’)

To seek further evidence for contribution of post-transcription to circadian whole-cell proteomes, we examined two properties of rhythmicity, amplitude and phase, in populations of genes in which both transcripts and whole-cell proteins are rhythmic under STD or ECD. In gene expression process, transcripts, which have an average half-life of 4.8 min(*35*), are produced before proteins. Therefore, a rhythmic gene population with more contribution from transcription would have higher ratio of genes with the phase of transcript leading the phase of protein to genes with the phase of transcript lagging the phase of protein. We found the ratio to be ∼4:1 under STD but ∼ 1:1 after ECD (Fig. 4H). Another prediction is that a rhythmic gene population with higher contribution from protein rhythmicity would have higher ratio of protein’s amplitude to transcript’s amplitude. The amplitude ratio is ∼1:4 under STD and increases to 9:1 after ECD (Fig. 4I). These results indicated that transcription contributes more to protein rhythmicity under STD than ECD while post-transcription contributes more to protein rhythmicity after ECD than STD. These are additional pieces of evidence for a higher contribution of post-transcription to protein rhythmicity after ECD.

### ECD reprograms circadian molecular functions

To assess the impact of ECD on circadian functions, we performed gene ontology enrichment analysis of rhythmic whole-cell proteins after ECD alone or in comparison with STD. Beside of known ECD-associated pathways such as C-type lectin receptor, Toll-like receptor or GnRH(*36–38*), the ECD rhythmic population is also enriched with post-transcriptional and post-translational processes such as mRNA processing, translation initiation, intracellular protein transport and MAPK/Ras signaling. Interestingly, there is an apparent day-night functional partitioning. Post-translational processes such as vesicular transport and protein localization peak during the nighttime while post-transcriptional and translational initiation processes peak during the daytime (Fig. 5A). In response to ECD, the enrichment of several STD-enriched processes is diminished such as acyl-CoA and di/tri-carboxylic acid metabolic processes while several others became highly enriched including post-transcription processing and intra-cellular protein transport. Among proteins undergoing significant change in rhythmicity are regulators of these processes, some of which were implicated in circadian dys/functions, such as PRMT5(*39*), SRSF6, XRCC5(*40*) (mRNA processing), RPLP0, EIF5, EIF4EBP2 (translation initiation), H/KRAS(*9*), AKT2(*41*) (Ras signaling), SEC61B, IPO9, EXOC6 (protein transport) (Figs. 5B-C). ECD thus reprograms a wide range of circadian functions at the molecular level.

**Fig. 5:**
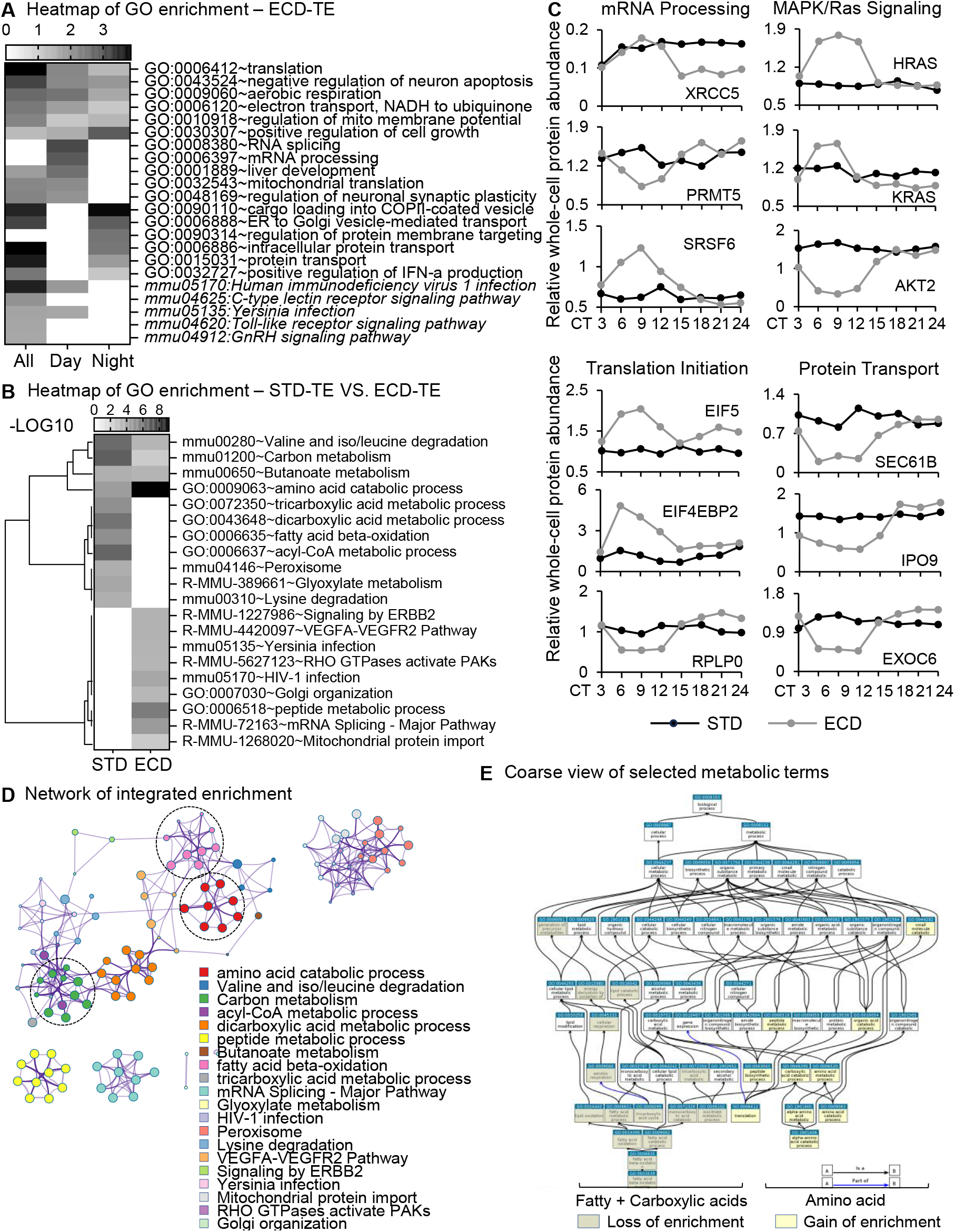
Comparative functional G.O. enrichment analyses. (**A-B**) Heatmap of G.O. enrichment analysis of (A) ECD whole-cell rhythmic proteins peaking during day-time, night-time or both, (B) whole-cell circadian proteome under ECD in comparison with STD. (**C**) Comparative whole-cell protein abundance throughout a circadian cycle of representative factors in mRNA processing, translation initiation, RTK/Ras signaling and protein transport under STD versus ECD conditions. (**D**) Reactome enrichment analysis of (B). (**E**) G.O. Slim map of changes in enrichment of terms related to fatty acid, carboxylic acid and amino acid, as circled in C, in response to ECD.

Given the central role of liver in metabolism, we took a closer look at the effect of ECD on metabolic functions. Analyzing the enrichment and network interaction of metabolic terms, we found that fatty acid and carboxylic acid metabolic processes lost most of their enrichment while amino acid metabolic processes, which were also enriched under STD, significantly increase in enrichment (Figs. 5D-E). These results suggest there is a change in prioritizing of circadian regulation from carbohydrate and fat to protein metabolism in response to ECD. The lost in timing of fat and carbohydrate metabolism might underlie, at least in part, ECD-associated metabolic disorders.

Interestingly, a previous study showed that mice, after being subjected to the same ECD and recovery paradigm, exhibited normal body temperature and running-wheel behavior under diurnal (LD) condition(*34*). Together with this observation, our results suggested organisms who experienced ECD might exhibit normal daily behavioral rhythms but not fully recover at the molecular level, at least after a short period of recovery time. The discordance in recovery at molecular and behavioral levels might indicate a diversion or a masking of behaviors by light. We favor the later scenario as these behaviorally “normal” mice are very susceptible to physiological challenges(*33, 34*).

In summary, our study showed that environmental circadian disrupted re-programs the circadian gene expression system and molecular circadian functions, which are not recovered, at least at the molecular level, after one week of recovery. Additionally, it also provided the first comprehensive documentation of changes in the circadian gene expression process in response to environmental circadian disruption, laying the foundation for deciphering the molecular underpinnings of shiftwork and jetlag.

## Acknowledgments

We thank Dr. Peter R. MacLeish and Dr. Carl H. Johnson for valuable comments and suggestions in the preparation of the manuscript. We also thank many colleagues who gave valuable comments and suggestions during our presentation of the study at the GRC-Chronobiology 2023.

## Funding

The study was supported by NIGMS grants 1 R16 GM146703-01 (HD) and SC1 GM135112-01A1 (KB).

HD is supported by NIGMS grant 1R16GM146703-01 and NEI grant R01EY026291

KB is supported by NIGMS grant SC1 GM135112-01A1

GT is supported by NEI grants R01EY026291 and R21EY031821

JPD is supported by NIGMS grant GM127044

AJD is supported by NIGMS grant R35GM136661

CE is supported by NIGMS grant GM127260

## Author contributions

HD incepted the project and analyzed the data.

HD designed the experiments with inputs from GT, AJD, JPD & MP.

HD performed the experiments with help from KB & CE.

HD wrote the manuscript with inputs from all authors.

## Competing interests

Authors declare that they have no competing interests.

## Data and materials availability

All data are available in the main text or the supplementary materials.

## Supplementary Materials

Materials and Methods

figs. S1 to S4

References (*42–46*)

## Notes

### Competing Interest Statement

The authors have declared no competing interest.

