## Supplemental materials for "Environmental circadian disruption re-programs liver circadian gene expression"

**The PDF file includes:**

Materials and Methods

Figs. S1 to S4

References

**Materials and methods**

Tissue collection: 48 wildtype (C57BL6/J) mice at around 12-week-old, purchased from Jackson Laboratory, were randomly divided into 16 groups of 3. Eight group were subjected to standard light-dark cycle (STD; ON at 6AM and OFF at 6PM) while the remaining groups were subjected to Environmental-Circadian Disruption light-dark regiment(*13*) (ECD; 4 cycles of 6-hr phase-advance light shift). After the last phase shift, ECD animals were recovered under STD condition for 1 week. All animals were then subjected to constant darkness (DD). Tissues were collected individually at 3-hr interval starting from the 2^nd^ day in DD. A small piece of each tissue was preserved in RNAlater solution (Invitrogen) and the rest was flashed frozen in liquid N_2_ for downstream RNA or protein analysis, respectively.

RNA-seq**:** RNA-seq analysis was performed by the Emory Genomic core for each of the 48 tissue samples using PolyA+ (mRNA) at 100 million reads (50M paired ends). Alignment to mm39 mouse genome was performed using STAR v2.5.2 with 76.2-88.1% uniquely aligned reads. After filtering out time series with abundance of less than 50 transcripts or detection in less than 32 of 48 samples, we successfully quantified 21166 transcript time series across both STD and ECD conditions.

Mass Spectrometry: Mass spectrometry analysis was performed by the Thermo Fisher Scientific Center for Multiplex Proteomics at Harvard Medical School on an Orbitrap Eclipse mass spectrometer. Whole-cell and nuclear extracts from each of the 48 livers were prepared independently as previously described(*42*, *43*). The triplicates were pooled, in equal mass, according to group and compartment. Approximately 50µg of each pooled sample was then subjected to 18-plexes, with 2 spike samples in each 18-plex, for parallel analysis. At 1% False Discovery Rate (FDR) of peptide spectral match and 5% FDR of protein identification, the analysis identified 8,294 proteins in extracts under ECA condition and 6,261 proteins in extracts after ECD condition. Of these identified proteins 5257 to 7314 protein time series were quantified for each of the 4 proteomes and were used for further analysis.

Circadian rhythmic detection**:** We used common denominator of two algorithms: RAIN and BIO_CYCLE. RAIN is a non-parametric algorithm that detects both symmetric and asymmetric waveforms with high reproducibility and recall while BIO_CYCLE is a parametric deep learning algorithm that could detect various waveforms with low false positive rate and high precision(*31*, *44*, *45*). We call an abundance time series significantly rhythmic only if its q (RAIN) and q(BIO_CYCLE) are both less than 0.05 for transcript or p(RAIN) and p(BIO_CYCLE) are both less than 0.05 for protein.

Gene Ontology and Cis-element enrichment analysis: G.O. analysis was performed using Metascape(*46*) with the background is all proteins that were quantified in this study. Cis-element enrichment analysis was performed using iRegulon algorithm(*28*) in 20kb upstream of the transcription start site.

Antibodies and RT-qPCR Probes

Antibodies: α-BMAL1 was a gift from Charles J. Weitz (Harvard Medical School); α-CLOCK (Abcam; ab3517); α-PER2 (Millipore; AB2202); α-NR1D1 (Abclonal; Catalog # A20452); α-SAP155 (Bethyl Laboratories, Inc; Catalog # A00-996A)

Probes:

1	mARNTL-qPCR-F	CAGAAGCAAACTACAAGCCAACA

2	mARNTL-qPCR-R	GGTCACATCCTACGACAAACA

3	mCLOCK-qPCR2-F	CCTTCAGCAGTCAGTCCATAAA

4	mCLOCK-qPCR2-R	CATGCCTTGTGGAATTGGTAAAT

5	mPER1-qPCR1-F	CCTGGAGGAATTGGAGCATATC

6	mPER1-qPCR1-R	CCTGCCTGCTCCGAAATATAG

7	mPER2-qPCR-F	CAACAACCCACACACCAAAC

8	mPER2-qPCR-R	CTCGATCAGATCCTGAGGTAGA

9	mPER3-qPCR-F	CACTCCAGGATGTGTGTTTCT

10	mPER3-qPCR-R	GATCTTCTGGGTGCAAGTATGT

11	mCRY1-qPCR-F	CTCAGTCCTTATCTCCGCTTTG

12	mCRY1-qPCR-R	CCACAGGAGTTGCCCATAAA

13	mCRY2-qPCR-F	GAGAACCATGACGACACCTATG

14	mCRY2-qPCR-R	AGCTTCTGTCTCTCCTCCTT

15	mCSNK1D-qPCR-F	TACTTCAACCTGGGCTCTCT

16	mCSNK1D-qPCR-R	CAATGGGAGTGGACATCTTCTT

17	mDbp-qPCR-F	CTGAGGAACAGAAGGATGAGAAG

18	mDbp-qPCR-R	TGGTTCTCCTTGAGTCTTCTTG

19	mLrwd1-qPCR-F	TGAAGAAGCTGAGGGAACTTG

20	mLrwd1-qPCR-R	TGTTGGTACAGCGAAGGATG

21	mCyc1-qPCR-F	ATTGCGAGAAGGCCTCTATTT

22	mCyc1-qPCR-R	TGCCATCATCATACTCCAAGAC

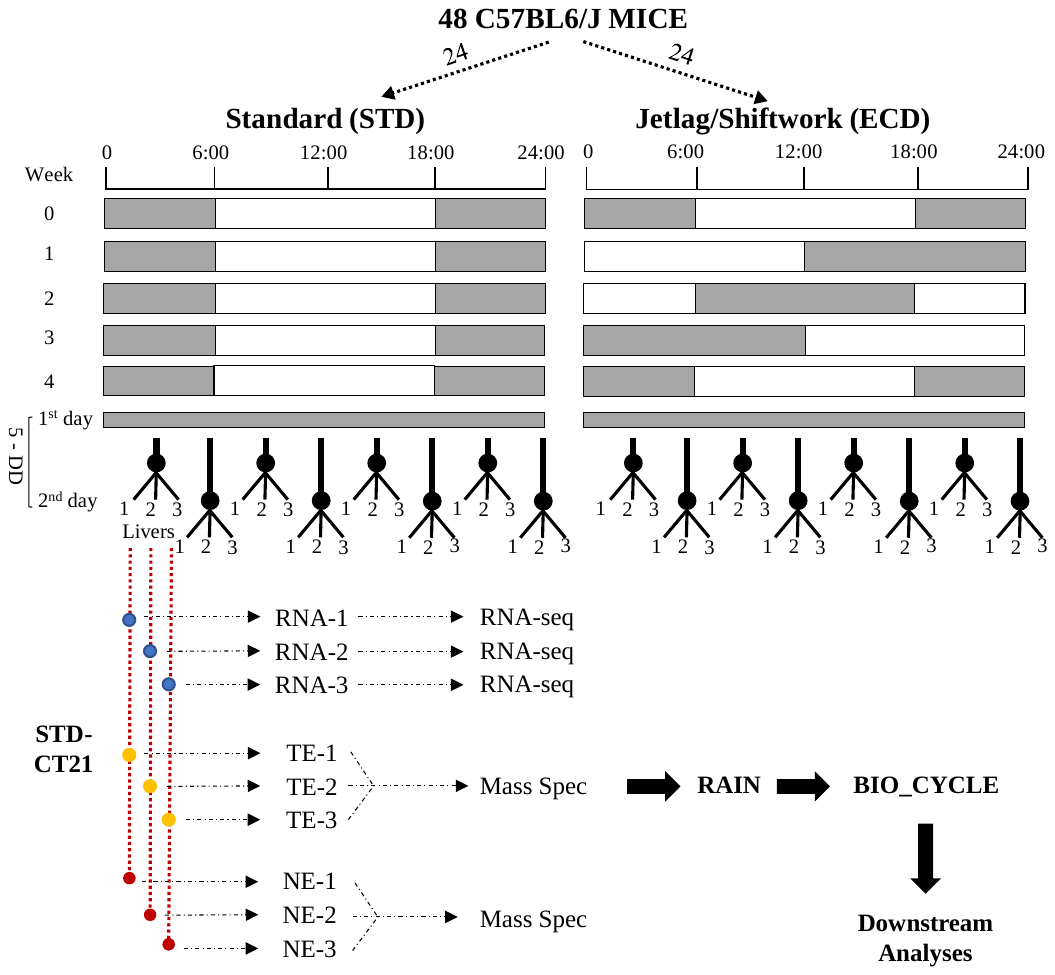

**fig. S1:** **Experimental paradigm**. 48 C57BL6/J mice of similar ages were divided into 16 groups of 3 mice/group. 8 groups were subjected to standard light-dark cycle (STD); ON at 6am and OFF at 6pm), while the remaining 8 groups were subjected to Jetlag/Shiftwork light-dark regiment (ECD; 4 cycles of 6-hr phase-advance light shift for 1 week per cycle). After the last shift, animals were recovered under the STD cycle for 1 week. All animals were then subjected to constant darkness (DD). Tissues from each group were collected under DD at 3-hr interval starting on the 2^nd^ day of the 5^th^ week. Each tissue was then processed for RNA extract, total extract (TE) and nuclear extract (NE), individually. RNA-seqs were performed for each RNA extract. For mass spectrometry, equal amount of the 3 extracts from the same group and compartment were pooled before the analysis. All time series of abundance were then subjected to both RAIN and BIO_CYCLE algorithms for determination of rhythmicity before proceeding to downstream analyses.

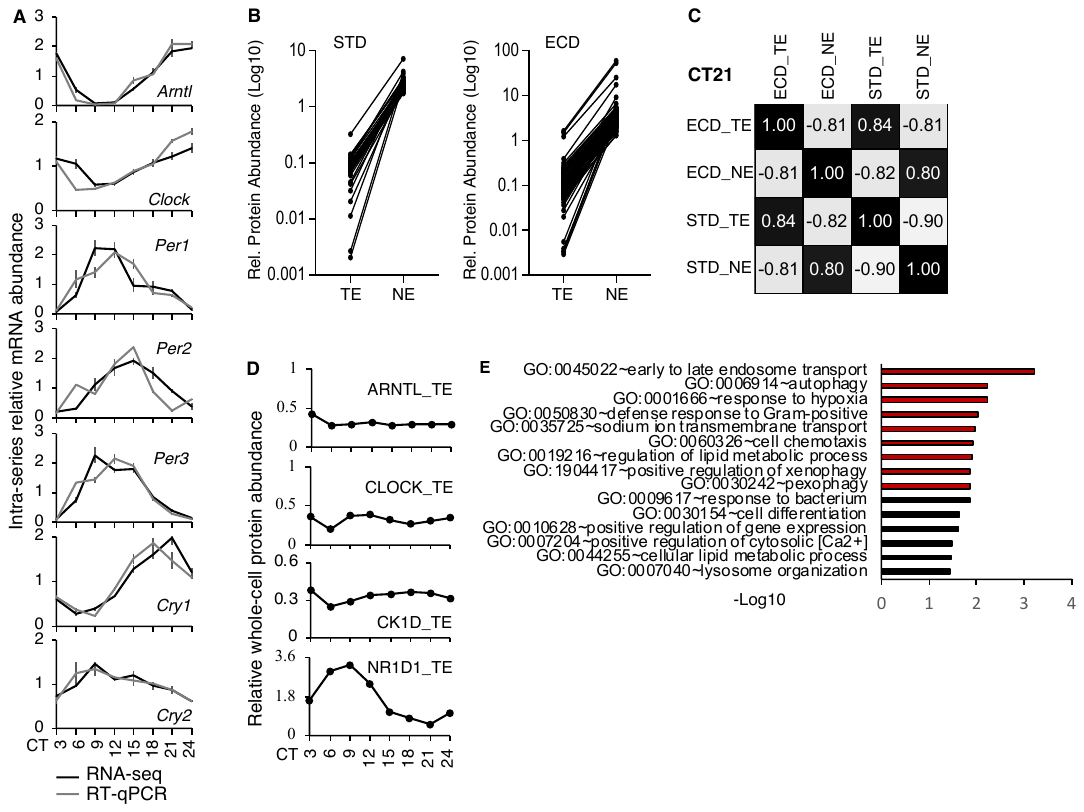

**fig. S2: Validations**. (**A**) Comparative abundance patterns of core clock transcripts detected by RNA-seq and RT-qPCR. (**B**) Series-average relative abundance in whole-cell and nuclear extracts of some known nuclear proteins with 20+ folds of NE/TE enrichment (Log10 scale). (**C**) Pearson’s correlation across compartments and conditions of all proteins quantified by mass spectrometry at CT21. (**D**) Patterns of whole-cell protein abundance of core clock components: ARNTL, CLOCK, CK1D and NR1D1 detected by mass spectrometry under STD. (**E**) G.O. enrichment analysis of the whole-cell circadian proteome under STD.

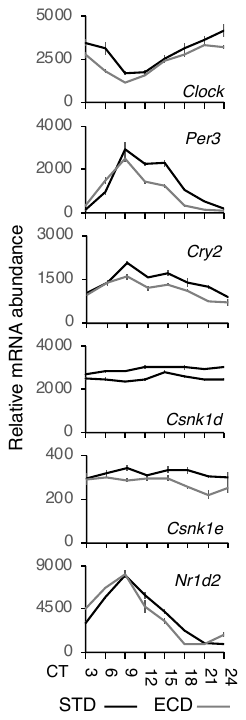

**fig. S3**: Core clock transcripts that showed no noticeable change in their circadian pattern in response to ECD as quantified by RNA-seqs.

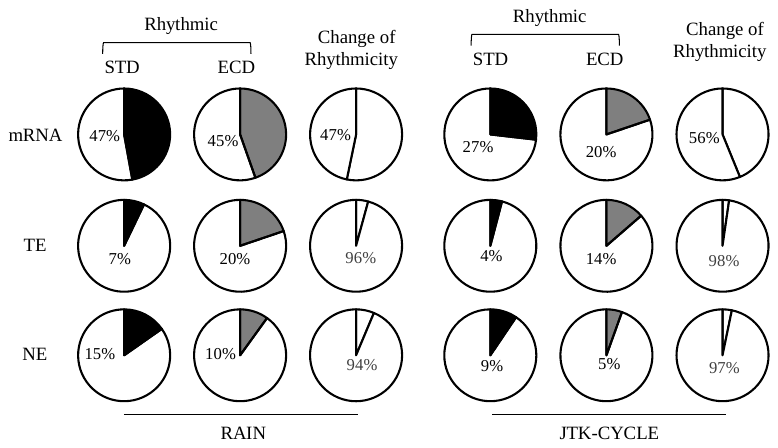

fig. S4. Rhythmic analysis using RAIN or JTK-CYCLE algorithm.
